# Class I histone deacetylases catalyze lysine lactylation

**DOI:** 10.1101/2025.02.25.640220

**Authors:** Takeshi Tsusaka, Mohd. Altaf Najar, Isha Sharma, Mariola M. Marcinkiewicz, Claudia Veronica Da Silva Crispim, Nathaniel W. Snyder, George M. Burslem, Emily L. Goldberg

**Affiliations:** Department of Physiology, University of California, San Francisco, San Francisco, CA 94158, USA; Department of Biochemistry and Biophysics, Perelman School of Medicine, University of Pennsylvania, Philadelphia, PA 19104 USA; Aging and Cardiovascular Discovery Center, Department of Cardiovascular Sciences, Lewis Katz School of Medicine, Temple University, Philadelphia, PA 19140, USA; Department of Cancer Biology and Epigenetics Institute, Perelman School of Medicine, University of Pennsylvania, Philadelphia, PA 19104 USA; Chan-Zuckerberg Biohub, San Francisco, CA 94158, USA

## Abstract

Metabolism and post-translational modifications (PTMs) are intrinsically linked and the number of identified metabolites that can covalently modify proteins continues to increase. This metabolism/PTM crosstalk is especially true for lactate, the product of anaerobic metabolism following glycolysis. Lactate forms an amide bond with the ε-amino group of lysine, a modification known as lysine lactylation, or Kla. Multiple independent mechanisms have been proposed in the formation of Kla, including p300/CBP-dependent transfer from lactyl-CoA, via a high-energy intermediate lactoylglutathione species that non-enzymatically lactylates proteins, and several enzymes are reported to have lactyl transferase capability. We recently discovered that class I histone deacetylases (HDACs) 1, 2, and 3 can all reverse their canonical chemical reaction to catalyze lysine β-hydroxybutyrylation. Here we tested the hypothesis that HDACs can also catalyze Kla formation. Using biochemical, pharmacological, and genetic approaches, we found that HDAC-catalyzed lysine lactylation accounts for the majority of Kla formation in cells. Dialysis experiments confirm this is a reversible reaction that depends on lactate concentration. We also directly quantified intracellular lactyl-CoA and found that Kla abundance can be uncoupled from lactyl-CoA levels. Therefore, we propose a model in which the majority of Kla is formed through enzymatic addition of lactate by HDACs 1, 2, and 3.

---

Modification of proteins by acyl groups, like acetylation, can control a range of protein functions, including location, structure, and association with binding partners (1). The overarching mechanisms controlling protein acylation are well studied: acyl groups can be added through enzymatic and non-enzymatic pathways and are removed enzymatically by deacylating enzymes like histone deacetylases (HDACs) and the sirtuins. While acetylation is the best studied protein acylation, other short-chain fatty acids like β-hydroxybutyrate (BHB) (2) and lactate (3) can also covalently modify lysine residues, although the physiological significance of these other acylations remains to be fully understood.

Lactate (l-Lactate) is a short-chain fatty acid produced and secreted by glycolytic cells (4). Lactate concentrations in blood and tissues can range from 0.5 – 20 mM and its production is acutely increased in response to a variety of stimuli, including intense exercise and sepsis (5-8). In addition to serving as a metabolic fuel, lactate can function as a signaling molecule. For example, lactate can be conjugated to phenylalanine to form N-lactoyl-phenylalanine (Lac-Phe) that gets secreted from cells and modifies food intake (9). Lactate can also form a covalent adduct on lysine residues, known as lactylation, that is found on histone and non-histone proteins, and has been reported in many cell types (3, 10-15).

Like other acylations, lysine l-lactylation (Kla) has been proposed to be formed through a lactyl-CoA-dependent pathway (3, 16). However, lactyl-CoA levels in cells are far less abundant compared to other CoA derivatives such as acetyl-CoA, propionyl-CoA, and succinyl-CoA (17). Interestingly, several enzymes have been reported to possess lactyl transferase activity and catalyze Kla formation on target proteins (18-21). HDAC6 was also recently reported to lactylate α-tubulin (22). Although less abundant than l-lactylation, the stereoisomer d-lactylation is formed from the non-enzymatic acyl transfer from the glycolytic intermediate S-D-lactoylglutathione (LGSH) generated in the glyoxylase pathway (23-25). Therefore, multiple non-redundant pathways may contribute to overall intracellular Kla levels.

We recently discovered that class I HDACs 1, 2, and 3 are reversible and catalyze lysine beta-hydoxybutyrylation (Kbhb) through a condensation reaction between the BHB carboxylic acid and free amine on lysine residues (26). In this new HDAC-catalyzed mechanism of protein acylation, we ruled out a requirement for an activated intermediate like BHB-CoA. We also showed that the same active site residues were required for both deacetylation and β-hydroxybutyrylation, consistent with our model that HDACs 1, 2, and 3 are reversible enzymes that catalyze the addition or removal of acyl chains. Of note, our model of HDAC-catalyzed protein acylation has been independently verified by others for lysine sorbylation (27, 28). As others have reported that HDACs 1, 2, and 3 can de-lactylate proteins (29), we hypothesized that, like Kbhb formation, HDACs also contribute to Kla formation through a similar reversible mechanism. Using biochemistry, molecular biology, and mass spectrometry, we confirmed HDACs 1, 2, and 3 are capable of directly catalyzing the formation of Kla in a recombinant reconstitution assay and in live cells. Our data further demonstrate that HDAC-catalyzed Kla formation is the dominant mechanism of protein lactylation.

## HDACs 1, 2, and 3 catalyze lysine lactylation

We previously used an in vitro reconstitution assay to show that recombinant HDAC2 (rHDAC2) can biochemically catalyze lysine acylation by short-chain fatty acids (26). We confirmed these results with lactate and verified that HDACs 1, 2, and 3 are each capable of this biochemical reaction and this could be inhibited by coincubation with the pan-HDAC active site inhibitor Trichostatin A (TSA) (Fig. 1A). We used liquid chromatography tandem mass spectrometry (LC-MS/MS) to validate the western blot data and identified histone-tail lysine residues K14, K18, K23, K27, and globular lysine K79 as lactylated sites by rHDAC2 in this in vitro assay (Fig. 1B-C). These data indicate that, like Kbhb, HDAC enzymatic activity is sufficient for Kla formation.

**Fig. 1.**
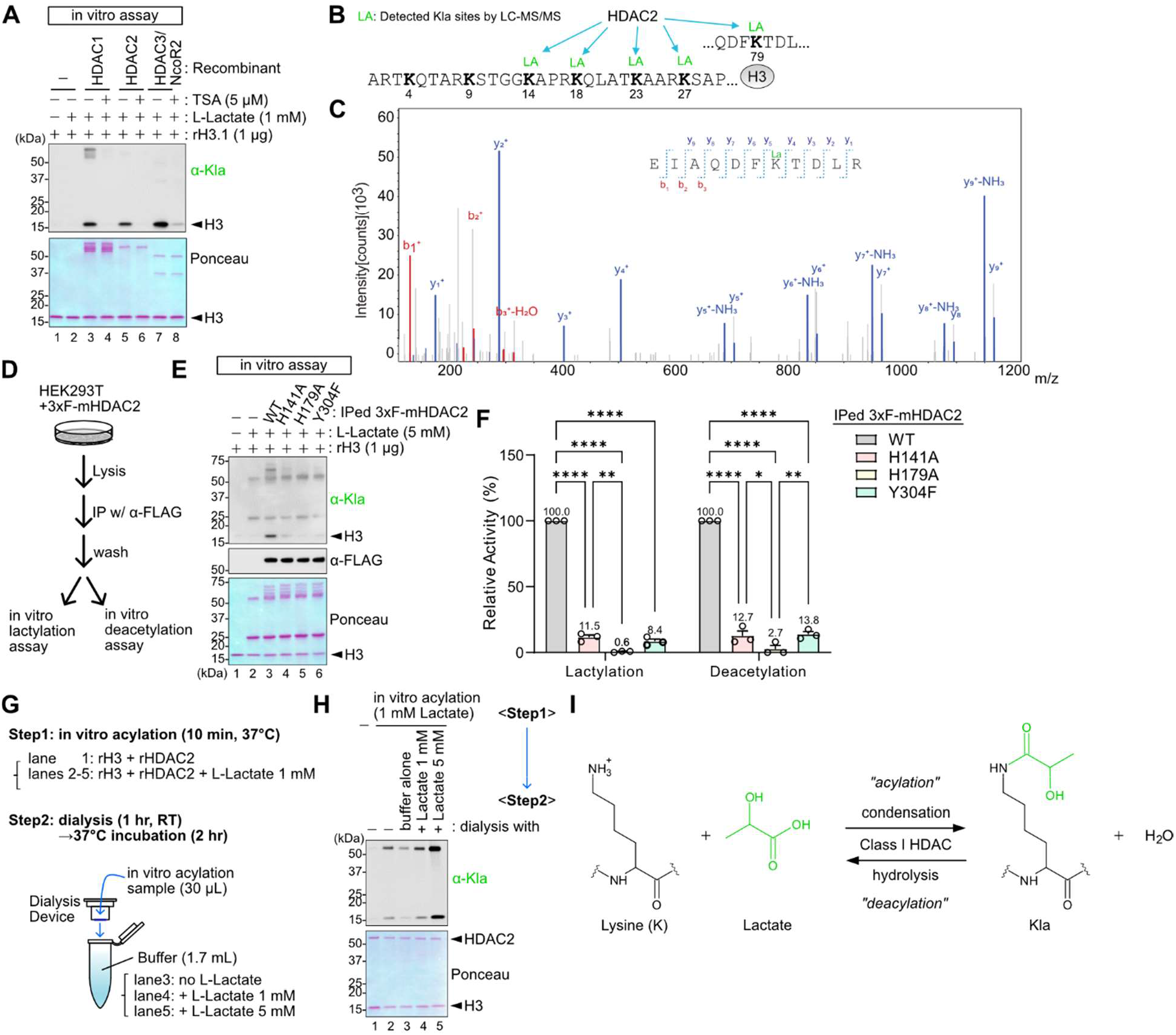
HDACs 1, 2, and 3 catalyze lysine lactylation in vitro. (**A**) Western blots of an in vitro lysine lactylation assay with recombinant HDAC1 (rHDAC1), rHDAC2, or rHDAC2/NcoR2 and histone H3 (rH3) in the presence of 1 mM L-Lactate with or without TSA (5 µM). The reactions were performed at 37°C for 30 mins. Protein loading was visualized by ponceau S staining, and Kbhb was detected by western blot. Experiments were repeated independently twice with similar results. (**B-C**) LC-MS/MS analysis of H3 after in vitro lysine lactylation assay. Detected Kla-modified sites on H3 (B). MS/MS-spectra for the indicated peptide with detected y and b ions (C). (**D**) Schematic of experimental workflow. HDAC2 KO HEK293T cells expressing 3xFLAG-mHDAC2 were used for immunoprecipitation with α-FLAG antibody. The immunoprecipitants were used for in vitro lysine lactylation assays and in vitro deacetylation assays. (**E**) Representative western blots of in vitro lysine lactylation with IPed 3xF-mHDAC2 WT or indicated mutants. This experiment was independently repeated three times with similar results. (**F**) Lysine lactylation and deacetylation activity for each mutant. Data are represented as mean +/-SEM activity relative to WT 3xF-mHDAC2, from n=3 independent experiments. (**G**) Schematic of the experimental workflow to test the Lactate concentration-dependence of HDAC reversibility using a dialysis system. (**H**) Representative western blot to assess Kla formation before and after dialysis with different L-Lactate concentrations. This experiment was repeated independently twice with similar results. (**I**) Schematic of the proposed model of the forward and reverse reactions catalyzed by class I HDACs.

We next tested the importance of key active site residues in lysine lactylation. We established HDAC2 knockout HEK293T cell lines expressing 3xFLAG-tagged mouse HDAC2 (mHDAC2) wild-type (WT) and individual point mutants H141A, H179A, and Y304F. These mutants each have a single substitution of residues required for deacetylation and lysine β-hydroxybutyrylation (26, 30). We purified these WT or mutant HDAC2 proteins using FLAG antibody-conjugated beads and performed our in vitro lactylation assay with rH3 as a model substrate. The WT 3xF-mHDAC2 showed the strongest lysine lactylation activity toward rH3, while all three mutants decreased the lactylation activity as expected (Fig. 1D-F). To test the reversibility of the enzymatic activity of rHDAC2 toward lysine lactylation, we used a dialysis system in combination with our in vitro lactylation assay to manipulate lactate concentrations (Fig. 1G-H). When lactate was removed from the reaction conditions by dialysis, rHDAC2 removed the lactyl modification from H3, consistent with a previous study (29). In contrast, lysine lactylation levels were retained or even increased when dialysis was performed in buffers containing the same or higher concentrations of lactate, respectively. Taken together, these data are consistent with a model in which the HDAC2 catalysis is reversible, enabling it to either add or remove lactate from lysine residues, dependent on the concentration of lactate (Fig 1I).

## HDAC-catalyzed lysine lactylation is the dominant source of Kla in cells

Next, we investigated the relevance of HDAC-catalyzed lysine lactylation in live cells. We treated HEK293T cells with several HDAC inhibitors (HDACi), including butyrate, TSA, SAHA, and MS-275. Of note, unlike Kbhb, which requires exogenous treatment of BHB in non-ketogenic cells, HEK293T cells basally contain many Kla-modified proteins, presumably due to their high metabolic and proliferative nature. All the HDAC inhibitors broadly decreased basal Kla levels and increased lysine acetylation (Kac) as expected (Fig. 2A-B, S1). Next, we knocked down HDAC1, HDAC2, and HDAC3 in varying combinations in HEK293T cells to further evaluate the role of HDACs in intracellular Kla formation. Each single knockdown showed a similar reduction (approximately ∼50%, lanes 3-5) in Kla levels, and the triple knockdown showed the strongest effect on Kla levels (lane 9), suggesting HDACs 1, 2, and 3 redundantly contribute to Kla formation in cells (Fig. 2C-D). We also confirmed that HDACi treatment did not affect intracellular lactate concentration (Fig. 2E), indicating that class I HDACs directly control Kla levels, rather than indirectly controlling lactate substrate availability to modify Kla abundance.

**Fig. 2.**
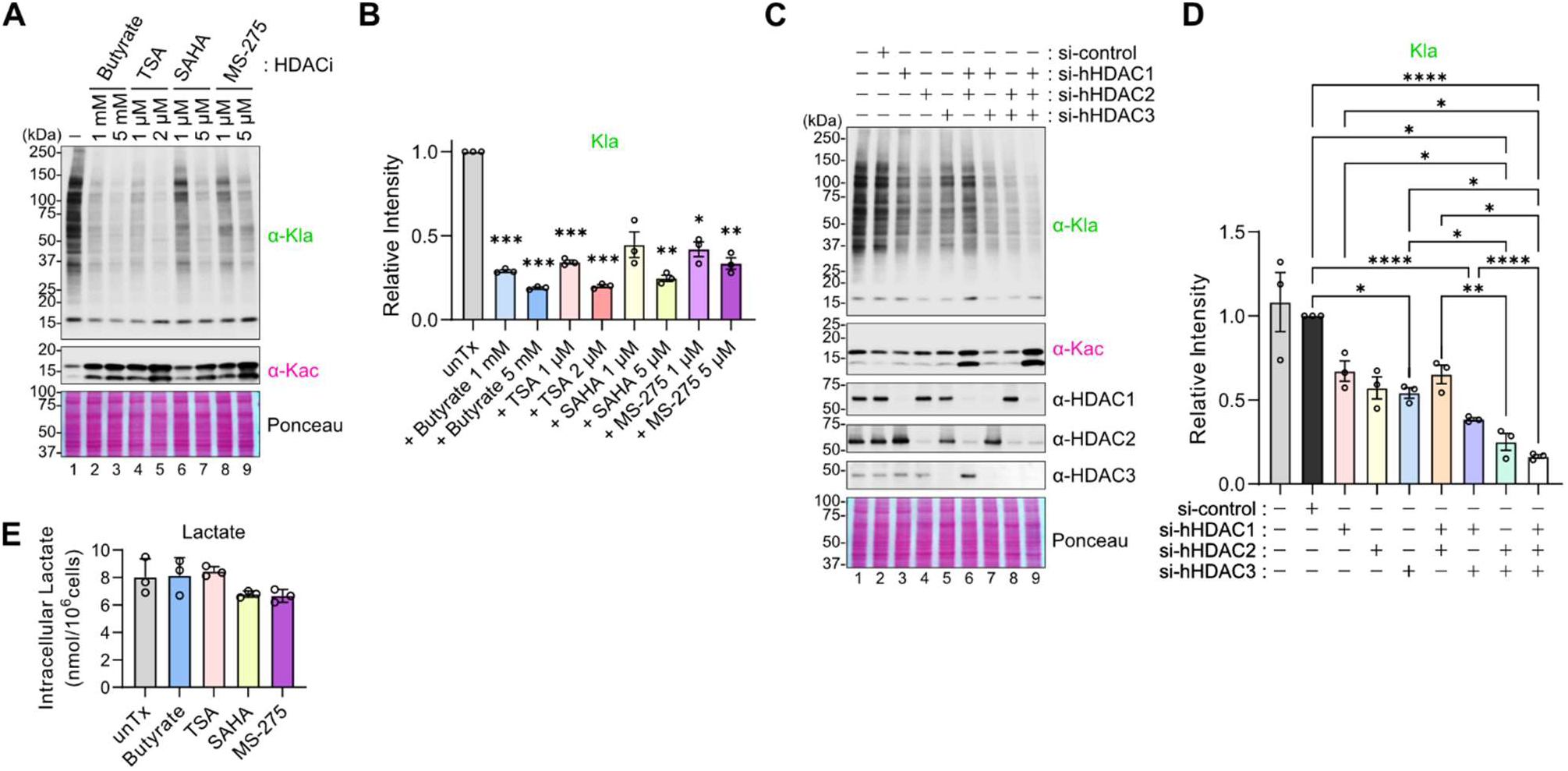
Catalytic activities of HDAC1, 2, and 3 are required for Kla formation in cells. (**A-B**) HEK293T cells were treated with each HDAC inhibitor at the indicated concentrations for 24 hours. Representative western blots are shown in (A). Quantification of anti-Kla signals relative to lane 1 (untreated, unTx) were shown in (B). Signals were normalized to ponceau S staining. Data are represented as mean +/-SEM of three independently performed experiments and each symbol represents an individual experiment. Statistical differences were calculated by 1-way ANOVA followed by Dunnett’s correction for multiple comparisons. (**C-D**) HEK293T cells were transfected with siRNA against the indicated targets. Representative western blots are shown in (C). Quantification of anti-Kla signals relative to lane 2 (si-control) are shown in (D). Signals were normalized by ponceau S staining. Data are represented as mean +/-SEM of three independently performed experiments and each symbol represents an individual experiment. Statistical differences were calculated by 1-way ANOVA followed by Tukey’s test for multiple comparisons. (**E**) Quantification of intracellular lactate concentrations in HEK293T cells treated with HDAC inhibitors: Butyrate at 5 mM, TSA at 1 µM, SAHA at 5 µM, MS-275 5 µM. Data are represented as mean +/-SD of technical triplicates. *p<0.05, **p<0.01, ***p<0.001, ****p<0.0001

## Kla formation is regulated by glucose metabolism

Because glycolysis-derived l-lactate and glyoxylase-derived d-lactate have both been implicated in Kla formation, we examined the importance of these isomers in HDAC-catalyzed Kla in cells. First, we generated LDHA/LDHB double knock-out (LDHA/B dKO) HEK293T cells to disrupt l-lactate production (Fig. S2, 3A). We observed a striking reduction in global Kla modifications in the LDHA/B dKO cells that was rescued by exogenous l-Lactate supplementation (Fig. 3B-C). We became aware of a preprint demonstrating that the pan-Kla antibody also detects carboxyethyl-lysine (Kce) modifications (31). Using their same in vitro system, we confirmed that at high in vitro concentrations, the pan-Kla antibody does react with Kce-modified BSA (Fig. S3A-B). Next, we knocked down GLO1 with siRNA, which would lead to the accumulation of methylglyoxal (MGO), and thus increase Kce and decrease d-lactyl-lysine (24, 25) but should have no impact on l-lactate or l-lactyl-lysine. GLO1 knockdown did not change global Kla levels as detected with the pan-anti-Kla antibody that recognizes l-lactyl-lysine (Fig. S3C-E). Moreover, when we used a pan-Kce antibody to detect Kce-modified proteins, in comparison to the Kce-modified BSA, only a very weak signal was detected in the cell extract samples, with no apparent change upon GLO1 knockdown, suggesting that this PTM might only be present at very low levels in cells. (Fig. S3F). Of note, we would expect our LDHA/B dKO to possibly also increase Kce modifications due to the buildup of pathway intermediates, but we observed no Kce-modified proteins in these cells either. These data strongly support our conclusion that the pan-Kla antibody is detecting lactylated proteins in our experiments and indicate that the lactate derives from glycolysis.

**Fig. 3.**
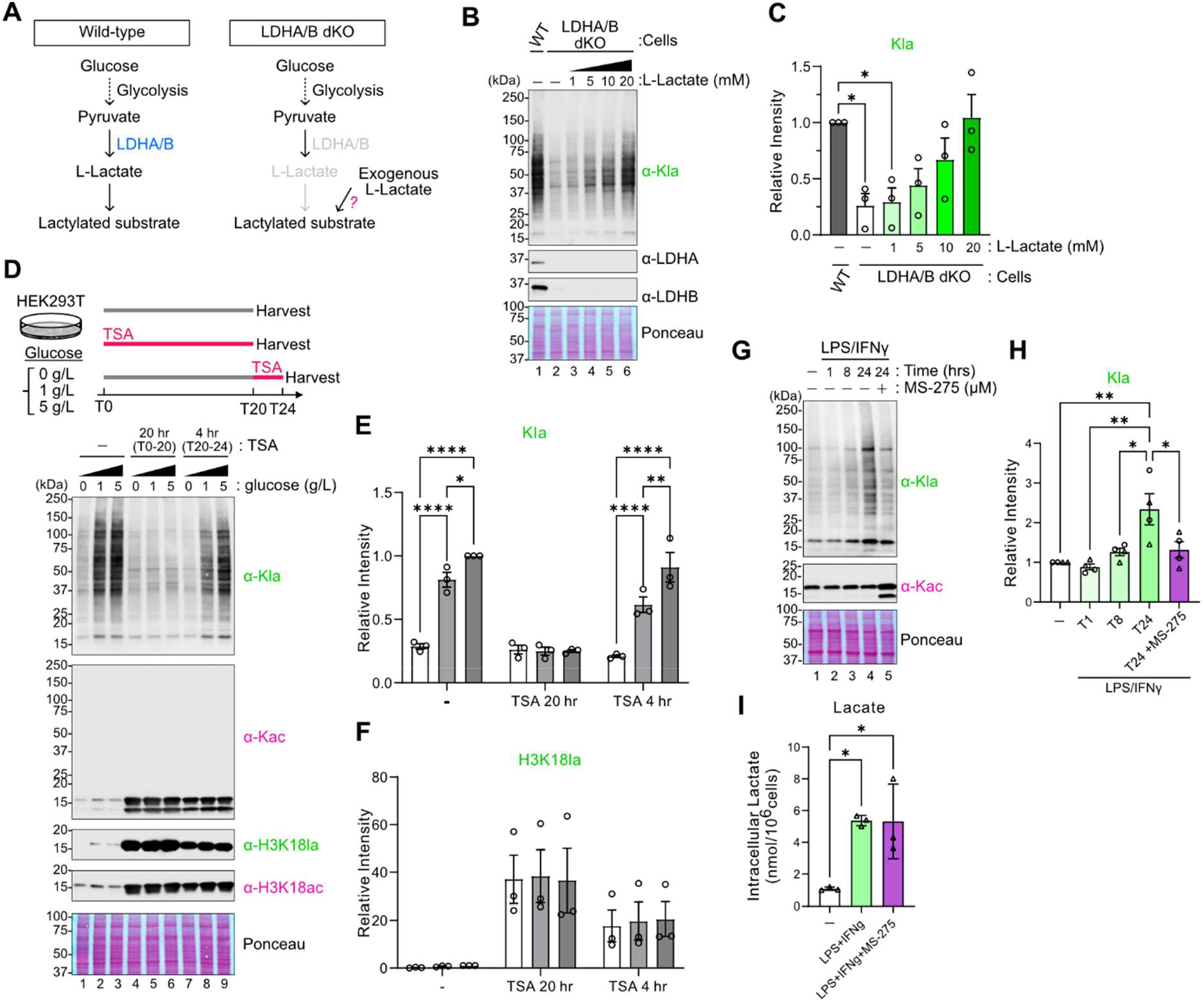
Kla formation is regulated by glucose metabolism. (**A**) Schematic of anticipated Kla levels in wild-type and established LDHA and LDHB double knock-out (LDHA/B dKO) cells. (**B-C**) LDHA/B dKO HEK293T cells were treated with L-Lactate at the indicated concentrations for 24 hours. Representative western blots are shown in (B) and quantification of anti-Kla signals relative to lane 1 (untreated, unTx) were shown in (C). Signals were normalized to ponceau S staining. Data are represented as mean +/-SEM of three independently performed experiments and each symbol represents an individual experiment. Statistical differences were calculated by 1-way ANOVA followed by Dunnett’s correction for multiple comparisons. (**D-F**) WT HEK293T cells were cultured in 0, 1, or 5 g/L glucose containing medium and treated with 1 µM TSA at the indicated time points. Representative western blots are shown in (D). Quantification of anti-Kla or anti-H3K18la signals relative to lane 3 (untreatment, 5 g/L glucose) were shown in (E) or (F), respectively. Signals were normalized to ponceau S staining. Data are represented as mean +/-SEM of three independently performed experiments and each symbol represents an individual experiment. Statistical differences were calculated by 1-way ANOVA followed by Tukey’s correction for multiple comparisons. (**G-I**) Bone-marrow derived macrophages (BMDM) were stimulated with LPS (5 ng/mL) and IFNγ (12 ng/mL) for the indicated time. MS-275 (0.5 µM) was added at 8 hr after LPS/IFNγ stimulation, resulting in 16 hours of inhibition from T8 to T24. Representative western blots are shown in (G). Quantification of anti-Kla signals relative to lane 1 (unstimulated) and normalized to ponceau S staining are shown in (H). Data are represented as mean +/-SEM of three independently performed experiments and each symbol represents an individual experiment. Statistical differences were calculated by 1-way ANOVA followed by Tukey’s correction for multiple comparisons. Quantification of intracellular lactate concentrations in BMDM after 24 hours of stimulation, with or without MS-275, are shown in (I). *p<0.05, **p<0.01, ****p<0.0001

Lactate is a byproduct of glycolysis and is secreted from highly glycolytic cells. As such, we cultured HEK293T cells under no, low (1 g/L), and high (5 g/L) glucose conditions for 24 hours and observed that Kla modification across all proteins correlated with glucose concentration and consequent lactate concentration (Fig. S4A-C). We also observed reduced global lysine lactylation in oxamate (OXA) or dichloroacetate (DCA)-treated cells (Fig. S4D-F), both of which can lower lactate production. We also confirmed that co-treating cells with TSA to inhibit HDAC enzymatic activity blocked the glucose-driven Kla accumulation (Fig. 3D, E), supporting our model that glycolysis-derived lactate is used in HDAC-catalyzed Kla formation. Since H3K18la was proposed to be involved in gene expression (3), modification at this position has gained significant attention, as evidenced by over 400 hits on Google Scholar as of February 2025. We therefore wanted to examine the contribution of HDACs on histone H3K18la using the commercial H3K18la antibody. The antibody used in the Zhang et al. has been discontinued so we tested the alternative anti-H3K18la from the same vendor (PTM-1406, PTM bio labs). We confirmed that, like global Kla, the H3K18la signals also decrease in cells deprived of glucose, as expected (Fig. 3D, F). However, the H3K18la signals in cells treated with TSA for 20 hrs exceeded those of untreated cells, regardless of glucose concentration (lanes 1-6). To rule out the possibility that residual lactate might be used for lysine lactylation, we also included the samples with the short exposure of TSA (4 hrs) following 20 hr glucose-restriction and obtained a similar result (lanes 7-9). Given that lactate and Kla-modified proteins are extremely limited in the absence of glucose, it is possible that the widely used H3K18la antibody might recognize other modification(s) in addition to lactylated lysine, similar to what we have shown for H3K9bhb (32) and is a well-known issue for histone modifications (33-35).

To further explore the metabolic regulation of Kla, we used a model of primary murine bone marrow-derived macrophages (BMDM). Activation of macrophages with the toll-like receptor 4 (TLR4) ligand lipopolysaccharide (LPS) leads to acute upregulation of glycolysis and lactate secretion. In agreement with a prior study (3), we confirmed that LPS or LPS+IFNγ activation also leads to Kla accumulation and importantly, this was blocked when BMDM were co-treated with the HDAC inhibitors MS-275 or butyrate during LPS treatment (Fig.3G-H, Fig. S3G-I). We also noted that LPS or LPS+IFNγ stimulation induced global lysine lactylation, not just on histones, and both of these were prevented by MS-275 or butyrate treatment. We also confirmed that MS-275 treatment did not affect lactate production in activated BMDM (Fig.3I). These results show that glycolytically activated macrophages rely on HDACs to promote Kla formation.

## HDAC-catalysis is the primary mechanism for Kla formation

Because multiple non-redundant pathways can promote Kla formation, we evaluated each of them individually for their role in global Kla levels. Recent studies found that the alanyl-tRNA synthetases, AARS1 and AARS2, act as a lactyl-transferase using lactate and ATP (19-21) (Fig. 4A). However, this function of AARS enzymes was studied in the context of glucose starvation or largely focused on lactylation of specific target proteins. To compare the relative contribution of AARS enzymes towards global basal levels of Kla in cells, we used siRNA to knockdown AARS1 and AARS2. However, the knockdown of either enzyme did not alter Kla levels (Fig. 4B-E), suggesting the AARS pathway is not a dominant source of global lysine lactylation. Another proposed model is lactyl-CoA dependent lysine lactylation (Fig. 4F). Two enzymes have been reported to be lactyl-CoA synthetases: ACSS2 (acetyl-CoA synthetase 2) and SUCLG2 (succinate-CoA ligase GDP-forming subunit beta) (36, 37). When we used siRNA to knockdown ACSS2 and SUCLG2 in HEK293T cells, we observed a modest but reproducible decrease in Kla after ACSS2 knockdown, but no effect after SUCLG2 knockdown (Fig. 4G-J). Next, we assessed the putative lactyl-transferase CBP and p300 and HDACs in l-Lactate-derived lysine lactylation. We tested if Kla rescue in lactate-supplemented LDHA/B-dKO cells was p300/CBP-dependent or HDAC-dependent. Importantly, when LDHA/B-dKO cells were treated with exogenous l-lactate to restore Kla levels, the p300/CBP inhibitor A485 did not prevent Kla formation, but HDAC1-3 inhibitor MS-275 still blocked Kla formation (Fig. 4K, L). Moreover, class I HDAC inhibition with MS-275 did not alter lactyl-CoA levels (Fig. 4M), uncoupling the lactyl-CoA and Kla levels and further emphasizing that a p300/CBP-dependent acyl transfer from lactyl-CoA is not the primary mechanism of Kla formation. These data support our model that HDAC-catalyzed Kla formation is the dominant mechanism of overall lysine lactylation.

**Fig. 4.**
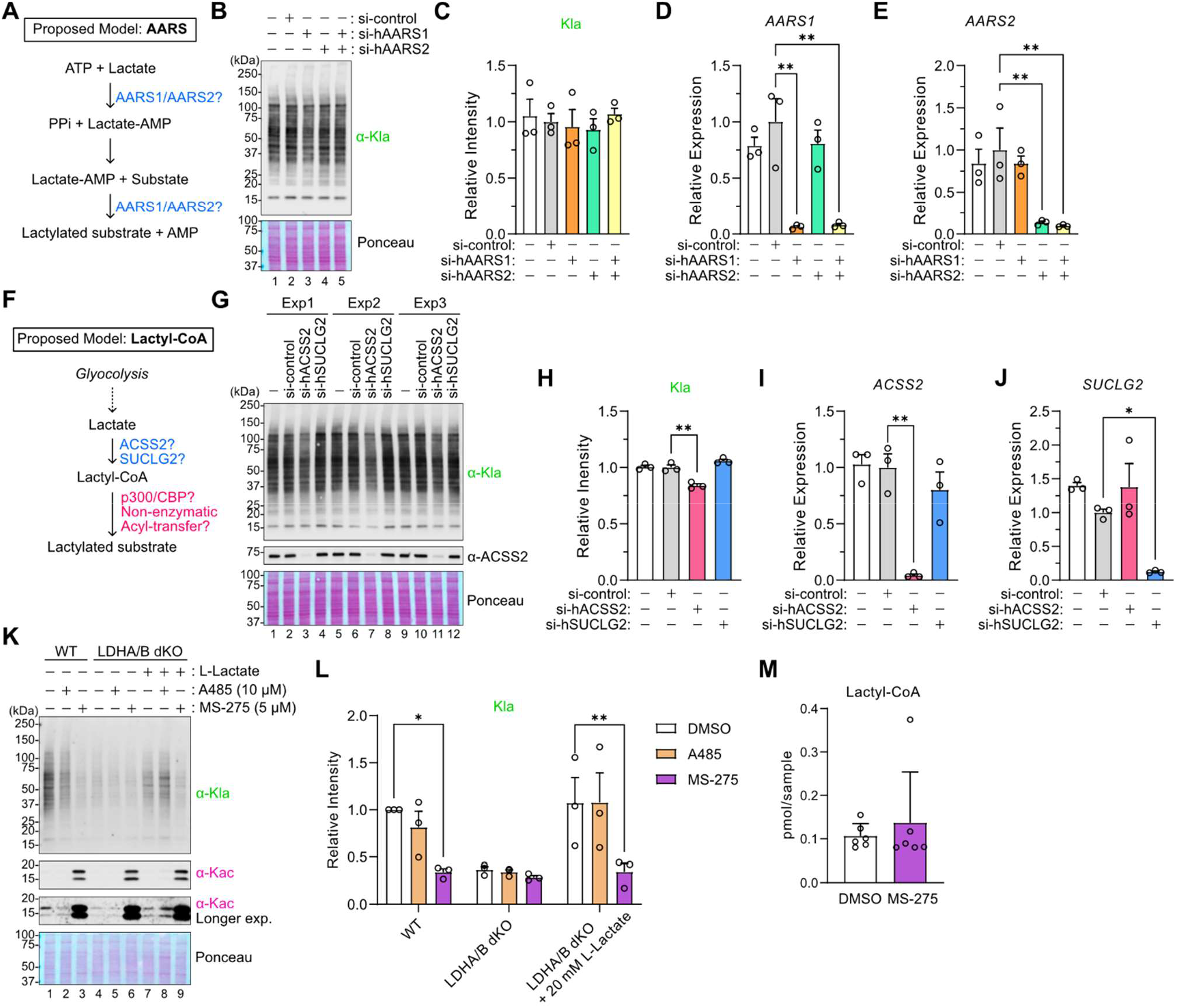
Lactate and HDAC enzymatic activity are the dominant regulators of Kla formation. (**A**) Proposed model of AARS enzymes-dependent lysine lactylation. (**B-E**) HEK293T cells were transfected with siRNA to assess the role of indicated target proteins in Kla formation. Representative western blots are shown in (B). Quantification of anti-Kla signals relative to lane 2 (si-control), normalized to ponceau S staining, are shown in (C). Relative mRNA expression for (D) AARS1, and (E) AARS2 were measured by qPCR and normalized to RPL13A. Data are represented as mean +/-SEM of three independently performed experiments and each symbol represents an individual experiment. Statistical differences were calculated by 1-way ANOVA followed by Dunnett’s correction for multiple comparisons. (**F**) Proposed model of lactyl-CoA enzyme-dependent lysine lactylation. (**G-J**) HEK293T cells were transfected with siRNA to assess the role of ACSS2 and SUCLG2 in Kla formation. Western blots containing all three independently performed experiments were shown in (G, labeled Exp1, Exp2, and Exp3). Quantification of anti-Kla signals relative to lane 2 (si-control) are shown in (H). Signals were normalized to ponceau S staining. Relative mRNA expression for (I) ACSS2 and (J) SUCLG2 were measured by qPCR and normalized to RPL13A. Data are represented as mean +/-SEM of three independently performed experiments and each symbol represents an individual experiment. Statistical differences were calculated by 1-way ANOVA followed by Dunnett’s correction for multiple comparisons. (**K-L**) LDHA/B dKO HEK293T cells were treated with 20 mM L-Lactate and A485/MS-275 inhibitors for 24 hours. Representative western blots are shown in (K) and quantification of anti-Kla signals relative to lane 1 (WT, untreated, unTx) are shown in (L). Signals were normalized to ponceau S staining. Data are represented as mean +/-SEM of three independently performed experiments and each symbol represents an individual experiment. Statistical differences were calculated by 1-way ANOVA followed by Dunnett’s correction for multiple comparisons. (**M**) Quantification of intracellular lactyl-CoA concentrations in HEK293T cells treated with MS-275 at 5 µM. Data are represented as mean +/-SD of 6 technical replicates. *p<0.05, **p<0.01, ***p<0.001, ****p<0.0001

## Discussion

Our data show that while lactyl-CoA may contribute to some Kla formation, global lysine lactylation is predominantly driven by class I HDACs and their enzymatic activity. This result is in agreement with our recent discovery that HDACs 1-3 catalyze Kbhb formation (26). We expect HDACs might utilize other short-chain fatty acids for known and yet-to-be-discovered lysine acylations. While the classical understanding of lysine acylation most often involves high-energy metabolites like acyl-CoAs that mediate acyl group transfer, we and others report independent alternative lysine lactylation mechanisms, including (i) the condensation reaction catalyzed by HDACs (ii), the transfer catalyzed by AARS using ATP and lactate (19-21), and (iii) non-enzymatic reaction via lactoyl-glutathione (24, 25). Future studies examining how these different mechanisms are balanced in different cell types or metabolic conditions, and how different target proteins may become modified, will enhance our collective understanding of protein acylation.

An open question remains as to the biological effects of HDAC-catalyzed lysine lactylation. We have uncovered some of the biochemical parameters of this pathway and confirmed that it operates in cells, but what purpose this mechanism accomplishes and under what conditions this pathway is activated remain unknown. During our studies, Li et al reported that HDAC6, a cytoplasmic HDAC that deacetylates tubulin, catalyzes the lactylation of tubulin via a condensation reaction (22). In addition, two preprints report that HDACs 1-3 catalyze lysine sorbylation (27, 28). Thus, our results are highly complementary and, together, provide a well-supported novel mechanism of protein acylation that is broadly relevant across HDAC family proteins that applies to multiple acylation substrates. Our study reveals the contribution of HDACs for lysine lactylation in cells and expands our understanding of how metabolically-sensitive modifications are established.

## Experimental procedures

### Mice

All mice were housed and maintained in a specific pathogen-free animal facility at the University of California, San Francisco. All animal protocols were approved by the UCSF Institutional Animal Care and Use Committee. Adult male and female mice (2-4 month old) were used for BMDM experiments. C57BL6/J mice were obtained from the Jackson Laboratory (stock #000664).

### Cell culture

HEK293T cells were generally cultured in high glucose (4.5 g/L ≈ 25 mM) DMEM (Gibco, 10569044) supplemented with 10% fetal bovine serum (FBS) and 1x antibiotic-antimycotic (Gibco, 15240062) in a 37°C incubator with 5% CO2. For glucose starvation/supplementation experiments, cells were cultured in no-glucose DMEM (Gibco, 11966025) supplemented with 10% dialyzed FBS (Gibco, A3382001), 1% GlutaMax (Gibco, 35050-061), 1 mM Sodium Pyruvate (Gibco, 11360-070), 1x antibiotic-antimycotic (Gibco, 15240062). D-glucose (G7021, Silgma-Aldrich) was dissolved in Ultrapure Distilled Water (10977015, Invitrogen) at 100 g/L as a stock, sterilized by filtration, and added to the medium to achieve the indicated glucose concentration. HDAC2 KO HEK293T cells and 3xFLAG-mHDAC2 wild-type (WT) or mutant-expressing cell lines were generated previously (26). Cells were treated as indicated in the respective figure legends with the following reagents: sodium L-Lactate (71718, Sigma-Aldrich), sodium butyrate (B5887, Sigma-Aldrich), Trichostatin A (TSA, T8552, Sigma-Aldrich), Suberoylanilide Hydroxamic Acid (SAHA, 10009929, Cayman), MS-275 (14043, Active Motif), A485 (63875, Tocris), sodium dichloroacetate (DCA, S8615, Selleckchem), sodium oxamate (OXA, S6871, Selleckchem).

Bone marrow-derived macrophages (BMDM) were established and cultured in RPMI supplemented with 10% FBS, 1x antibiotic-antimycotic, and 20 ng/mL MCSF (Peprotech #315-02), for 7 days, as described previously (38). For stimulation, BMDMs were seeded at 1×106 cells/ml in a 24-well plate. After an over-night incubation, pre-warmed (37°C) medium containing 6x stimulants was added to the culture to achieve a final concentration of LPS (O11:B4,1 µg/mL) or LPS (5 ng/mL) + IFNγ (12 ng/mL, eBioscience #14-8311-63).

### siRNA transfection

Cells were transfected with siRNA as described previously (26). Briefly, siRNA at 10 nM, Opti-MEM, and RNAiMax (Life Technology) were mixed in each well of a 6-well dish. After a 10 min incubation, cell suspensions (4×105 cells/well) were added to each well, and antibiotic-free medium was added to bring the culture volume to 2 mL. After 48 hours, cells were harvested, and the siRNA transfection was repeated once more.

The next day, the medium was replaced with pre-warmed (37°C) fresh medium, and after at least 4 hours of incubation, cells were collected for western blot or qPCR analysis. The following Silencer Select siRNA (Thermo Fisher Scientific) were used: s73 (HDAC1, validated), s6495 (HDAC2, validated), s16876 (HDAC3, validated), s842 (AARS1, validated), s33175 and s33177 (AARS2), s31745 and s31746 (ACSS2), s16775 and s16776 (SUCLG2), and s5825 (GLO1, validated). Efficient knock-downs for each gene were validated by qPCR or western blot analysis. For AARS2, ACSS2, and SUCLG2, two siRNAs were mixed and used for efficient knockdown and to avoid off-target effects.

### Plasmids

Oligo DNA that contained each gRNA sequence encoding human LDHA or LDHB was inserted into lentiGuide-Puro vector (52963, Addgene) according to the Addgene resource information. The following gRNA sequences were used: g-hLDHA#1 (ACAACTGTAATCTTATTCTG), g-hLDHA#2 (AGCCGTGATAATGACCAGCT), g-hLDHA#4 (GGGGAACATGGAGATTCCAG), g-hLDHB#1 (TACATCCACTTCCAATCACG), g-hLDHB#2 (GGACTGTACTTGACGATCTG). All plasmids used in this study were confirmed by Sanger sequencing.

### CRISPR/Cas9-mediated gene knockout in cells

Cas9-expressing HEK293T cells were previously established (26). Cas9+ HEK293T cells were reverse-transfected with each gRNA-expressing vector using Lipofectamine 3000 and selected with puromycin at 2 µg/mL and blastcidin at 10 µg/mL. After confirming the efficient reduction of target protein in bulk cells by western blot, single clones for each gRNA were cloned and tested. For LDHA KO, we isolated 5 clones and used the LDHA KO#1 cl.2 for g-hLDHB transfection. For LDHA/B dKO, we isolated 5 clones and used the LDHA/B dKO #2 cl.4 for Fig. 3-4.

### Western blot analysis

Western blot analysis was performed as described previously (32). Briefly, cells were harvested after trypsinization and quenching with medium, washed once with PBS, and lysed with RIPA buffer (Thermo Scientific) supplemented with protease inhibitors (Thermo Scientific). The cell lysates were sonicated using a Branson Sonifier 450 until the viscosity of lysates disappeared and then centrifuged at 14,000 rpm (17,968 xg) at 4°C for 5 mins to clarify cell lysates. Protein was quantified with the DC Protein Assay Kit (BioRad Laboratories). SDS-PAGE samples were prepared by mixing clarified lysates with 4x Laemmli buffer/10% β-mercaptoethanol (BME) (BioRad Laboratories) and then boiled at 95°C for 5 mins. Equivalent amounts of protein were resolved by SDS-PAGE and then transferred to a nitrocellulose membrane using the Trans-Blot Turbo Transfer system (BioRad Laboratories). Membranes were stained with Ponceaus S, blocked with 5% milk/TBS-T (Tris Buffered Saline/0.1% Tween-20) for 30-60 mins, and incubated with primary antibodies, followed by secondary HRP-conjugated antibodies. Blots were developed using a chemiluminescent substrate (SuperSignal™ West Pico PLUS or SuperSignal™ West Femto Maximum Sensitivity Substrate; Thermo Scientific) and imaged with an Azure 300 (Azure Biosystems). Ponceau S staining was used to confirm equal total protein loading across samples.

### Antibodies

The following antibodies were used in this study: anti-Kac antibody mix (1:1000-3000, CST, 9814S), anti-Kla (1:1000, PTM Biolabs, PTM-1401RM), anti-Kce (1:1000, PTM Biolabs, PTM-1701RM), anti-H3K18ac (1:3000, Abcam, ab1191), anti-H3K18la (1:1000, PTM Biolabs, PTM-1406), anti-HDAC1 (1:1000, CST, 5356), anti-HDAC2 (1:1000, CST, 5113), anti-HDAC3 (1:500, CST, 3949), anti-FLAG (1:3000, Sigma-aldrich, F3165), anti-ACSS2 (1:1000, CST, 3658), Goat anti-Rabbit IgG HRP (1:5000, Thermo Scientific, 31460), Goat anti-Mouse IgG HRP (1:5000, Thermo Scientific, 62-6520).

### RNA extraction and qPCR analysis

Total RNA was isolated using the RNeasy micro kit (Qiagen, 74034) according to the manufacturer’s protocol. cDNA was generated from equal amounts of RNA using iScript cDNA synthesis kit (Bio-rad, 1708891). Quantitative PCR (qPCR) with Power SYBR™ Green PCR Master Mix (Applied Biosystems™, 4367659) was carried out using CFX384 real-time PCR detection system (Bio-rad). The primer sequences are provided in Supplementary Table 1.

### In vitro lactylation assay with recombinant HDACs

In vitro lactylation assays were performed as described previously (26). Most in vitro reconstitution experiments used recombinant proteins from Cayman: rHDAC1 (10009231), rHDAC2 (36419), rHDAC3/NCOR2 (10009232). rHDAC2 from BPS Bioscience (50002) was used for Fig. 1B. We selected vendors based on the availability of recombinant proteins from each company at that time and observed no difference in rHDAC2 acylation capability among the vendors. Recombinant Histone H3.1 (rH3) was from NEB (M2503S). In most experiments, 1 µg rHDAC and/or 1 µg rHDAC was incubated in reaction buffer (50 mM Tris-HCl pH 7.5) with sodium L-lactate (71718, Sigma-Aldrich) for the indicated time at 37°C. Reactions were performed in 8-strip tubes in a thermal cycler (ProFlex PCR system, applied biosystems) in a final reaction volume of 15 µL. Reactions were stopped by adding 4x Laemmli buffer and subsequent boiling at 95°C for 5 mins or acetic acid (final. 0.15%) for western blot or mass-spectrometric analysis, respectively.

### LC-MS/MS analysis for in vitro acylated histone proteins

#### Derivatization/propionylation

Propionylation of lysine residue was performed using propionic anhydride and isopropanol in a 1:3 ratio. Equal amounts of protein between conditions were adjusted to pH 8 using NH4OH before the addition of 10 uL of propionylation reagents to each sample followed by vortexing, pH was again adjusted to 8 by adding 5-8 µl of NH4OH. The reaction was incubated at 37 °C for 15 minutes and repeated a total of 3 times to ensure completed propionylation.

#### Trypsin Proteolysis

Propionylated histones were diluted with 10 µl of 50 mM ammonium bicarbonate buffer and digested using Trypsin at a 1:50 ratio of protein to trypsin, overnight at at 37 °C. Extracts were combined, concentrated to dryness by SpeedVac, reconstituted in 0.1 % formic acid and desalted using C18 Stage-Tips. Digested peptides were dried using SpeedVac, dried peptides were cleaned using C18 Stage-Tips before LC-MS/MS.

#### LC-MS/MS analysis

The C18 cleaned peptides was analyzed on Thermo Scientific Orbitrap Exploris 240 mass spectrometer interfaced with Thermo Scientific UltiMate 3000 HPLC and UHPLC Systems. Peptide digests were reconstituted in 0.1 % formic acid and were separated on an analytical column (75 µm × 15 cm) at a flow rate of 300 nL/min using a step gradient of 1– 25 % solvent B (0.1 % formic acid in 100 % acetonitrile) for the first 50 minutes and 25–30 % for next 2 minutes, 30-70 % for 2 minutes, 70–1 % for next 2 minutes the total run time was set to 60 min. The mass spectrometer was operated in data-dependent acquisition mode. A survey full scan MS (from m/z 400–1600) was acquired in the Orbitrap with a resolution of 6000. Data were acquired in topN with 13 dependent scans. Fragmented using normalized collision energy with 37 % and detected at a mass resolution of 60,000. Dynamic exclusion was set for 8 s with a 10 ppm mass window.

#### Data Analysis

MS/MS searches were carried out using SEQUEST search algorithms against a custom-made database for human histones using Proteome Discoverer (Version 3.0, Thermo Fisher Scientific, Bremen, Germany). The workflow included Spectrum files, Spectrum selector, SEQUEST search nodes, Target decoy PSM validator, IMP-ptmRS, peptide validator, event detector, precursor quantifier. Oxidation of methionine, propionylation at lysine, lactylation at lysine and N-terminal protein acetylation were used as dynamic modifications. MS and MS/MS mass tolerances were set to 10 ppm and 0.02 Da, respectively. Trypsin was specified as protease and a maximum of two missed cleavage was allowed. Target-decoy database searches used for calculation of false discovery rate (FDR) and for peptide identification FDR was set at 1 %. Feature mapper and precursor ion quantifier were used for label-free quantification.

### In vitro deacetylation assay

In vitro deacetylation assays were performed as described previously(26). FLAG-IPed HDAC (WT and mutants) proteins were transferred to a 96-well half area assay plate (3994, CORNING) and the buffer was completely removed. Deacetylation assay with fluorescent molecule-conjugated peptide was performed using HDAC Fluorometric Activity Assay Kit (10011563, Cayman), according to the manufacture’s protocol. The reaction volume was 85 µL and deacetylation reaction was performed for 30 mins. The fluorescent signals in duplicates were detected by plate reader at excitation=350nm and emission=460nm (SpectraMax iD5, Molecular Devices). Relative IP 3xFLAG-mHDAC2 amounts were estimated based on in-parallel anti-FLAG western blot analysis and used for the normalization toward both in vitro lysine lactylation activity and in vitro deacetylation activity.

### Dialysis following in vitro lactylation experiment

In vitro lacylation was performed as described in “In vitro lactylation assay with recombinant HDACs”, with 2 µg rHDAC, 2 µg rHDAC, and/or 1 mM L-Lactate for 10 mins at 37°C, performed in 8-strip tubes in a thermal cycler (ProFlex PCR system, Applied Biosystems) in a final reaction volume of 30 µL. After the in vitro lactylation, four samples (lanes 2-5) were pooled, and 30 uL of the mixture was transferred to Slide-A-Lyzer™ MINI Dialysis Devices (3.5K MWCO, 69550, Thermo Scientific). The dialysis devices were submerged in 1.7 mL of 50 mM Tris-HCl pH7.5 buffer, containing either no L-Lactate, 1 mM L-Lactate, or 5 mM L-Lactate, and incubated for 1 hour at room temperature (23°C) with continuous agitation at 300 rpm using a heat block (Thermo Scientific). For a control sample (lane 2), 30 uL mixture within the dialysis device was left for 1 hour on a 2 mL tube without any solution, and the reaction was stopped by adding 4x Laemmli buffer and subsequent boiling at 95°C for 5 mins. For dialysis samples (lanes 3-5), after dialysis at room temperature, samples were transferred to 8-strip tubes and further incubated at 37°C for 2 hours to facilitate enzymatic reactions. Reactions were stopped by adding 4x Laemmli buffer and subsequent boiling at 95°C for 5 mins for western blot.

### Lactate quantification

Cells were harvested after trypsinization and quenching with their respective culture supernatants to maintain extracellular lactate concentrations during quenching and cell counting. After counting, cells were washed once with PBS, resuspended with 25 µL metaphosphoric acid (MPA) per 1.0×10^6 cells and incubated on ice for 5 mins. Following centrifugation at 10,000 xg for 5 mins at 4c, 20 µL of the supernatant was transferred to a new tube and neutralized with 1 µL of potassium carbonate to bring pH to approximately 8.5. The samples were diluted the sample 1:2 with 1x Assay buffer (50 mM potassium phosphate, pH 7.5), and lactate quantification assay was performed using L-Lactate Assay Kit (700510, Cayman) based on LDH catalysis, according to the manufacture’s protocol. The reaction volume was 100 µL, and LDH reaction was performed for 20 mins. The fluorescent signals in duplicates were detected by plate reader at excitation = 535 nm and emission = 587 nm (SpectraMax iD5, Molecular Devices). Intracellular lactate levels were quantified based on a lactate standard curve and normalized to cell numbers.

### Sample preparation for Lactyl-CoA quantification

For HDAC inhibitor-treated samples, cells were seeded at 2.5×106 cells/well in a 10-cm dish. After one day of culture, MS-275 (5 µM) or DMSO was directly added to the culture medium. After 24 hours, the culture medium was removed, directly resuspended in 1 mL of 10% (w/v) trichloroacetic acid (TCA), and stored at -80 °C until further processing. Six dishes were used as technical replicates for each condition.

### Liquid Chromatography Mass Spectrometry detection of lactyl-CoA

Acyl-CoAs were analyzed by liquid chromatography-high-resolution mass spectrometry (LC-HRMS) as previously described (17). 50 µL of short-chain acyl-CoA ISTD was added and then cell suspensions were sonicated with 5 x 0.5-second pulses at 50% intensity (Fisherbrand™ Sonic Dismembrator Model 120 with Qsonica CL-18 sonicator probe). Lysates were centrifuged at 17000 x g for 10 minutes at 4°C and clarified lysates were transferred to a deep-well 96-well plate for loading in a Tomtec Quadra4 liquid handling workstation. On the liquid handling workstation, lysates were applied to an Oasis HLB 96-well elution plate (30 mg of sorbent per well) pre-conditioned and equilibrated with 1 mL of methanol and 1 mL of water, respectively. After de-salting with 1 mL of water, acetyl-CoA was eluted into a deep-well 96-well plate using 1 mL of 25 mM ammonium acetate in methanol. Eluent was evaporated dried under nitrogen gas. A 12 point calibration curve of lactyl-CoA from 0.024 to 25 pmol/sample was prepared along with the samples. The dried LC-HRMS samples were resuspended in 50 µL of 5% (w/v) sulfosalicylic acid in water. 5 µL injections of each sample were analyzed via LC-HRMS. Samples were analyzed using an Vanquish Duo ultra-high performance liquid chromatograph coupled with a Q Exactive Plus mass spectrometer (Thermo Scientific) as previously described. A modified gradient using solvent A (5mM ammonium acetate in water), solvent B (5 mM ammonium acetate in 95:5 (v:v) acetonitrile: water) and solvent C (0.1% (v/v) formic acid in 80:20 (v:v) acetonitrile: water). Data was acquired using XCalibur 4.0 (Thermo Scientific) and analyzed using Tracefinder 5.1 (Thermo Scientific).

### Non-enzymatic in vitro carboxy-ethyllation with MGO

Bovine serum albumin (BSA, A9647, Sigma-Aldrich) was incubated in 20 mM phosphate buffer (pH 8.5) with or without methylglyoxal (MGO, 1 mM, M0252, Sigma-Aldrich) at 37°C over-night.

### Statistical analyses

Statistical analyses were performed in GraphPad Prism (version 10) unless otherwise indicated. We used two-tailed unpaired t-tests to calculate statistical differences between two groups. In analyses in which we compared multiple test conditions to a control group, we used 1-way analysis of variance (ANOVA) with post-hoc two-sided Dunnett’s test to calculate statistical differences. For multiple comparisons between all groups, we used 1-way ANOVA with Tukey’s post-hoc test to calculate statistical differences. Analysis information for each figure is indicated in the respective figure legends. For all experiments, p <0.05 was considered significant. *p<0.05, **p<0.01, ***p<0.001, ****p<0.0001.

## Data availability

All data related to this manuscript are contained within the main text and supplementary figures.

## Supporting information

This article contains supporting information.

Supplementary Figures S1-4.

Supplementary Table 1 for oligonucleotide sequences used for qPCR analysis.

## Acknowledgements

We thank past and present members of the Goldberg lab for their comments and intellectual discussion. We thank Hiten Madhani and Danica Fujimori for helpful intellectual discussion. We also thank Anna Molofsky’s lab (UCSF) for sharing HEK293T cells. We thank Jordan Meier (NCI) for the synthesis of lactyl-CoA standard used in LC-MS quantification.

## Funding and additional information

The Goldberg lab is funded by the Chan Zuckerberg Biohub (to ELG), the Sandler Program for Breakthrough Biomedical Research, which is partially funded by the Sandler Foundation (to ELG), and JSPS Overseas Research Fellowships (to TT). The Burslem lab is funded by NIGMS (R35GM142505) to GMB. Snyder Lab contributions to this project were funded by R01DK138011 to NWS.

## Author Contributions

ELG and TT conceptualized the project, designed experiments, and prepared the manuscript. TT performed most of the experiments and analyzed data. IS performed LDHA/B-dKO and Kce experiments under the supervision of TT and ELG. MAN performed LC-MS/MS sample prep, acquisition, and analysis of in vitro reconstitution samples under the supervision of GMB. MMM and CVDSC performed lactyl-CoA quantifications under the supervision of NWS. All authors assisted in manuscript preparation.

## References

1. Verdin, E., and Ott, M. (2015) 50 years of protein acetylation: from gene regulation to epigenetics, metabolism and beyond Nat Rev Mol Cell Biol 16, 258–264

2. Xie, Z., Zhang, D., Chung, D., Tang, Z., Huang, H., Dai, L. et al. (2016) Metabolic Regulation of Gene Expression by Histone Lysine beta-Hydroxybutyrylation Molecular cell 62, 194–206

3. Zhang, D., Tang, Z., Huang, H., Zhou, G., Cui, C., Weng, Y. et al. (2019) Metabolic regulation of gene expression by histone lactylation Nature 574, 575–580

4. Vander Heiden, M. G., Cantley, L. C., and Thompson, C. B. (2009) Understanding the Warburg effect: the metabolic requirements of cell proliferation Science 324, 1029–1033

5. Olenchock, B. A., Rathmell, J. C., and Vander Heiden, M. G. (2017) Biochemical Underpinnings of Immune Cell Metabolic Phenotypes Immunity 46, 703–713

6. Kelly, B., and O’Neill, L. A. (2015) Metabolic reprogramming in macrophages and dendritic cells in innate immunity Cell Res 25, 771–784

7. Van Hall, G., Jensen-Urstad, M., Rosdahl, H., Holmberg, H. C., Saltin, B., and Calbet, J. A. (2003) Leg and arm lactate and substrate kinetics during exercise Am J Physiol Endocrinol Metab 284, E193–205

8. Lewis, G. D., Farrell, L., Wood, M. J., Martinovic, M., Arany, Z., Rowe, G. C. et al. (2010) Metabolic signatures of exercise in human plasma Sci Transl Med 2, 33ra37

9. Li, V. L., He, Y., Contrepois, K., Liu, H., Kim, J. T., Wiggenhorn, A. L. et al. (2022) An exercise-inducible metabolite that suppresses feeding and obesity Nature 606, 785–790

10. Reynolds, M. B., Klein, B., McFadden, M. J., Judge, N. K., Navarrete, H. E., Michmerhuizen, B. C. et al. (2024) Type I interferon governs immunometabolic checkpoints that coordinate inflammation during Staphylococcal infection Cell Rep 43, 114607

11. Dichtl, S., Lindenthal, L., Zeitler, L., Behnke, K., Schlosser, D., Strobl, B. et al. (2021) Lactate and IL6 define separable paths of inflammatory metabolic adaptation Sci Adv 7,

12. Yang, Z., Yan, C., Ma, J., Peng, P., Ren, X., Cai, S. et al. (2023) Lactylome analysis suggests lactylation-dependent mechanisms of metabolic adaptation in hepatocellular carcinoma Nat Metab 5, 61–79

13. Zhang, N., Zhang, Y., Xu, J., Wang, P., Wu, B., Lu, S. et al. (2023) alpha-myosin heavy chain lactylation maintains sarcomeric structure and function and alleviates the development of heart failure Cell Res 33, 679–698

14. Zhang, X., Liu, Y., and Wang, N. (2025) Dynamic changes in histone lysine lactylation during meiosis prophase I in mouse spermatogenesis Proc Natl Acad Sci U S A 122, e2418693122

15. Yang, Z., Zheng, Y., and Gao, Q. (2024) Lysine lactylation in the regulation of tumor biology Trends Endocrinol Metab 35, 720–731

16. Zhang, D., Gao, J., Zhu, Z., Mao, Q., Xu, Z., Singh, P. K. et al. (2025) Lysine L-lactylation is the dominant lactylation isomer induced by glycolysis Nat Chem Biol 21, 91–99

17. Varner, E. L., Trefely, S., Bartee, D., von Krusenstiern, E., Izzo, L., Bekeova, C. et al. (2020) Quantification of lactoyl-CoA (lactyl-CoA) by liquid chromatography mass spectrometry in mammalian cells and tissues Open Biol 10, 200187

18. Niu, Z., Chen, C., Wang, S., Lu, C., Wu, Z., Wang, A. et al. (2024) HBO1 catalyzes lysine lactylation and mediates histone H3K9la to regulate gene transcription Nat Commun 15, 3561

19. Zong, Z., Xie, F., Wang, S., Wu, X., Zhang, Z., Yang, B. et al. (2024) Alanyl-tRNA synthetase, AARS1, is a lactate sensor and lactyltransferase that lactylates p53 and contributes to tumorigenesis Cell 187, 2375–2392 e2333

20. Ju, J., Zhang, H., Lin, M., Yan, Z., An, L., Cao, Z. et al. (2024) The alanyl-tRNA synthetase AARS1 moonlights as a lactyltransferase to promote YAP signaling in gastric cancer J Clin Invest 134,

21. Li, H., Liu, C., Li, R., Zhou, L., Ran, Y., Yang, Q. et al. (2024) AARS1 and AARS2 sense L-lactate to regulate cGAS as global lysine lactyltransferases Nature 634, 1229–1237

22. Sun, S., Xu, Z., He, L., Shen, Y., Yan, Y., Lv, X. et al. (2024) Metabolic regulation of cytoskeleton functions by HDAC6-catalyzed alpha-tubulin lactylation Nat Commun 15, 8377

23. Trujillo, M. N., Jennings, E. Q., Hoffman, E. A., Zhang, H., Phoebe, A. M., Mastin, G. E. et al. (2024) Lactoylglutathione promotes inflammatory signaling in macrophages through histone lactoylation Mol Metab 81, 101888

24. Gaffney, D. O., Jennings, E. Q., Anderson, C. C., Marentette, J. O., Shi, T., Schou Oxvig, A. M. et al. (2020) Non-enzymatic Lysine Lactoylation of Glycolytic Enzymes Cell Chem Biol 27, 206–213 e206

25. Zhao, Q., Wang, Q., Yao, Q., Yang, Z., Li, W., Cheng, X. et al. (2025) Nonenzymatic lysine D-lactylation induced by glyoxalase II substrate SLG dampens inflammatory immune responses Cell Res 35, 97–116

26. Tsusaka, T., Najar, M. A., Schwarz, B., Bohrnsen, E., Oses-Prieto, J. A., Lee, C. et al. (2023) Reversible histone deacetylase activity catalyzes lysine acylation bioRxiv 10.1101/2023.11.17.5675492023.2011.2017.567549

27. Sin, Y.-C., Abernathy, B., Mashek, D. G., and Chen, Y. (2024) Sorbate Induces Lysine Sorbylation Through Non-Canonical Activities of Class I HDACs to Regulate the Expression of Inflammation Genes bioRxiv 10.1101/2024.12.11.6279732024.2012.2011.627973

28. Kota Noritsugu, Y. S., Takehiro Suzuki, Komei Aoki, Tamiko Nozaki, Shigehiro A. Kawashima, Masaya Yamazaki, Kazuya Yamagata, Kota Koike, Kosuke Dodo, Kenji Ogawa, Mikiko Sodeoka, Minoru Yoshida, Naoshi Dohmae, Yoshito Kumagai, Akihiro Ito (2005) HDAC1 and HDAC2 are bidirectional enzymes that catalyze histone sorbylation to induce epigenetic alterations Research Square (Preprint) 10.21203/rs.3.rs-5847857/v1

29. Moreno-Yruela, C., Zhang, D., Wei, W., Baek, M., Liu, W., Gao, J. et al. (2022) Class I histone deacetylases (HDAC1-3) are histone lysine delactylases Sci Adv 8, eabi6696

30. Finnin, M. S., Donigian, J. R., Cohen, A., Richon, V. M., Rifkind, R. A., Marks, P. A. et al. (1999) Structures of a histone deacetylase homologue bound to the TSA and SAHA inhibitors Nature 401, 188–193

31. Kulkarni, C. A., and Brookes, P. (2020) Many Routes from Glycolysis to Histone PTMs. Nature “Matters Arising” response to: Zhang et al. Metabolic regulation of gene expression by histone lactylation. (2019) Nature. 574 (7779), 575–580. 10.31219/osf.io/sba8j

32. Tsusaka, T., Oses-Prieto, J. A., Lee, C., DeFelice, B. C., Burlingame, A. L., and Goldberg, E. L. (2023) Non-specific recognition of histone modifications by H3K9bhb antibody iScience 26, 107235

33. Rothbart, S. B., Dickson, B. M., Raab, J. R., Grzybowski, A. T., Krajewski, K., Guo, A. H. et al. (2015) An Interactive Database for the Assessment of Histone Antibody Specificity Molecular cell 59, 502–511

34. Fuchs, S. M., and Strahl, B. D. (2011) Antibody recognition of histone post-translational modifications: emerging issues and future prospects Epigenomics 3, 247–249

35. Egelhofer, T. A., Minoda, A., Klugman, S., Lee, K., Kolasinska-Zwierz, P., Alekseyenko, A. A. et al. (2011) An assessment of histone-modification antibody quality Nat Struct Mol Biol 18, 91–93

36. Zhu, R., Ye, X., Lu, X., Xiao, L., Yuan, M., Zhao, H. et al. (2025) ACSS2 acts as a lactyl-CoA synthetase and couples KAT2A to function as a lactyltransferase for histone lactylation and tumor immune evasion Cell Metab 37, 361–376 e367

37. Liu, R., Ren, X., Park, Y. E., Feng, H., Sheng, X., Song, X. et al. (2025) Nuclear GTPSCS functions as a lactyl-CoA synthetase to promote histone lactylation and gliomagenesis Cell Metab 37, 377–394 e379

38. Goldberg, E. L., Letian, A., Dlugos, T., Leveau, C., and Dixit, V. D. (2023) Innate immune cell-intrinsic ketogenesis is dispensable for organismal metabolism and age-related inflammation J Biol Chem 299, 103005

