## Supplemental figures for "Class I histone deacetylases catalyze lysine lactylation"

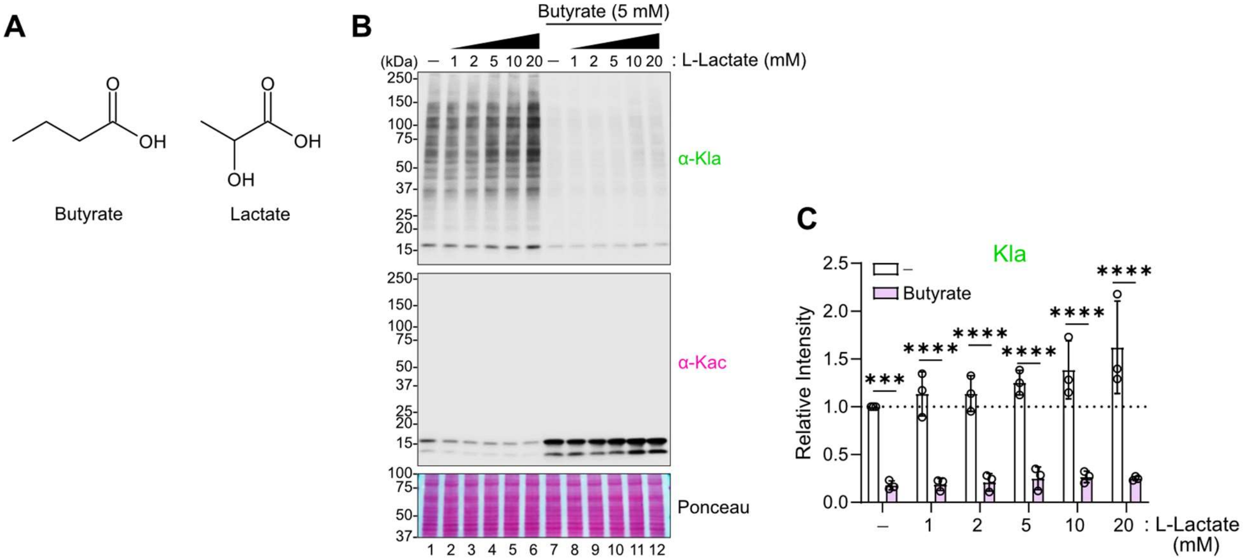


**Figure S1. Butyrate, a known histone deacetylation inhibitor, reduces Kla levels**

(**A**) Chemical structures of similar carboxylic acids, butyrate and lactate. (**B-C**) HEK293T cells were treated with L-Lactate and/or butyrate at the indicated concentrations for 24 hours. Representative western blots were shown in (B). Quantification of anti-Kla signals relative to lane 1 (untreatment, unTx) are shown in (C). Signals were normalized to ponceau S staining. Data are represented as mean +/- SEM of three independently performed experiments and each symbol represents an individual experiment. Statistical differences were calculated by Sidak's multiple comparisons test to compare with or without butyrate treatment. ***$p<0.001$, ****$p<0.0001$

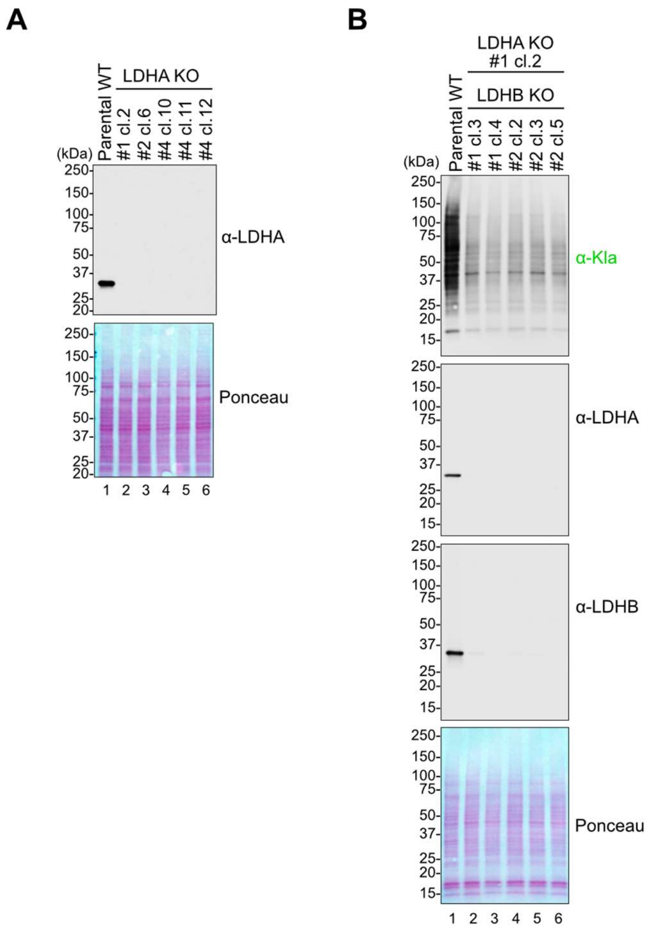


**Figure S2. Establishment of LDHA/B dKO HEK293T cells**

(**A**) Cas9-expressing HEK293T cells were transfected with gRNA targeting for *LDHA* gene. After isolating individual clones, the knock-out of LDHA in each clone was verified by western blot analysis. (**B**) The #1 cl.2 LDHA KO cells were transfected with gRNA targeting for *LDHB* gene. After isolating some clones, the knock-out of LDHB in each clone were verified by western blot analysis. The LDHA/B dKO #1 cl.4 cell line was chosen for subsequent analyses.

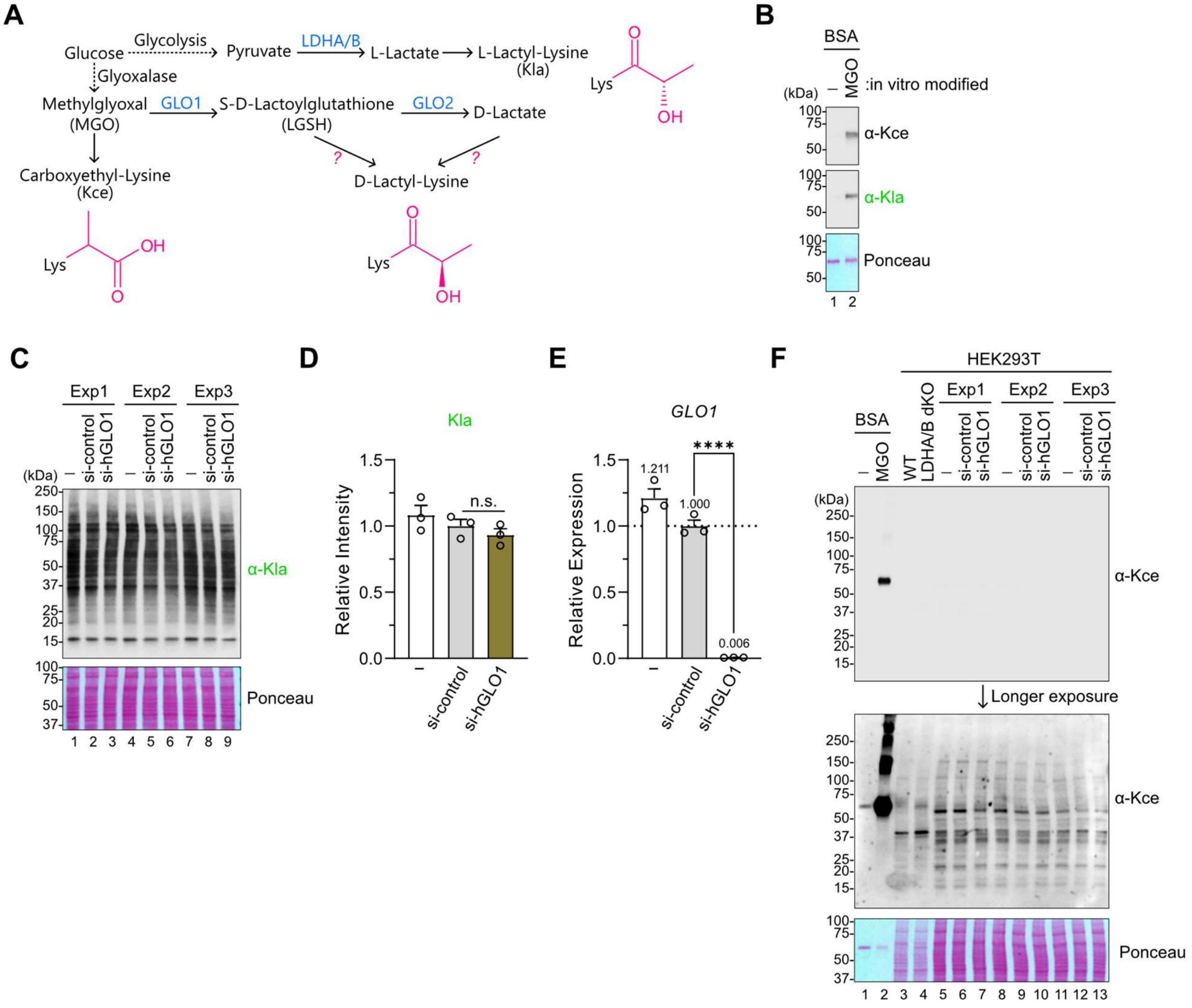


**Figure S3. Kla antibody detection is not confounded by intracellular Kce modification**

(**A**) Schematic of metabolic pathways that could produce lysine L-Lactylation (Kla), lysine D-lactylation, and lysine carboxyethylation. (**B**) Western blot analysis of BSA (Bovine serum albumin) incubated with MGO (methylglyoxal) that spontaneously produces Kce modification. (**C-F**) HEK293T cells were transfected with siRNA to assess the role of GLO1 and its associated Kce and D-lactyl-lysine in anti-Kla recognition. Western blots containing all three independently performed experiments are shown in (C, labeled as Exp1, Exp2, and Exp3). Quantification of anti-Kla signals relative to lane 2 (si-control), normalized to ponceau S staining, are shown in (D). Relative mRNA expression for (E) *GLO1*, was measured by qPCR analysis and normalized to *RPL13A* expression. Data are represented as mean +/- SEM of three independently performed experiments and each symbol represents an individual experiment. Statistical differences were calculated by unpaired t-test. ****p<0.0001, n.s. not significant. Western blots of the same samples as in (C) along with positive and negative control samples for Kla and Kce are shown in (F). We used the same samples of WT and LDHA/B dKO samples used in Fig. 3B (lanes 1-2). The BSA in vitro modified with MGO was freshly prepared.

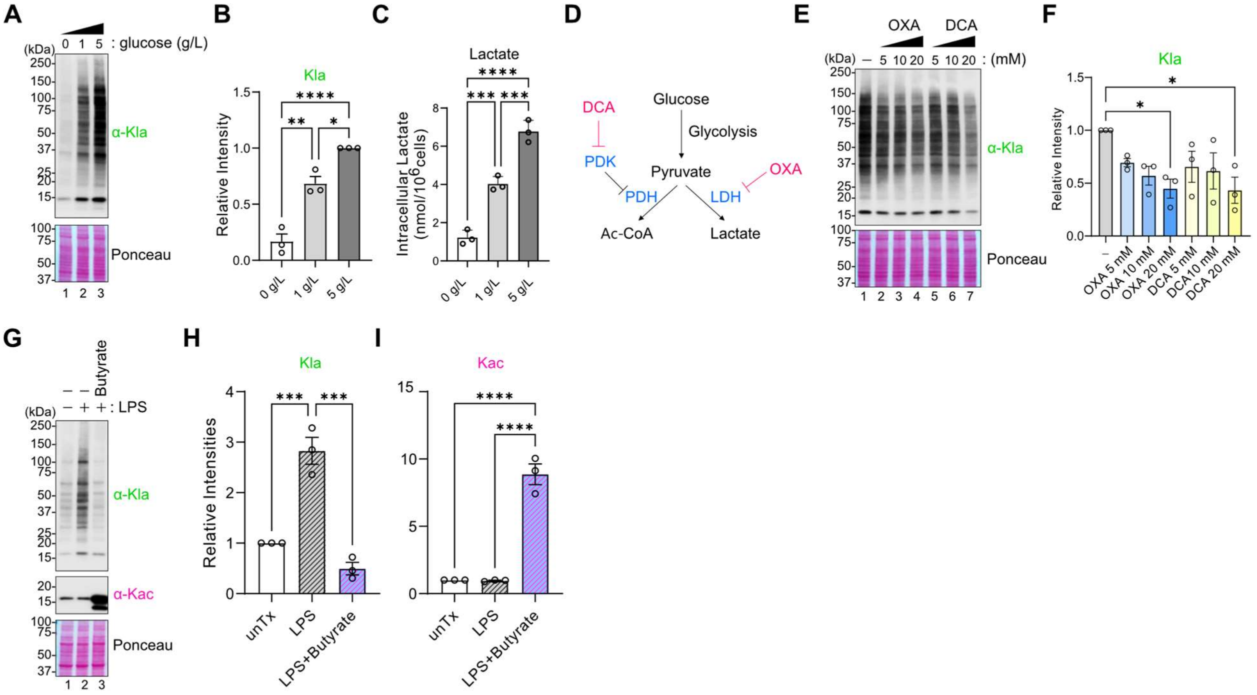


**Figure S4. Kla formation is regulated by glucose metabolism**

(**A-C**) WT HEK293T cells were cultured in 0, 1, or 5 g/L glucose-containing medium for 24 hours. Representative western blots are shown in (A). Quantification of anti-Kla relative to lane 3 (5 g/L glucose) is shown in (B). Signals were normalized to ponceau S staining. Data are represented as mean +/- SEM of three independently performed experiments and each symbol represents an individual experiment. Quantification of intracellular lactate concentrations in HEK293T cells are shown in (C). Data are represented as mean +/- SD of technical triplicates. Statistical differences were calculated by 1-way ANOVA followed by Tukey's correction for multiple comparisons. (**D**) Schematic of metabolic pathways and inhibitors. DCA (Dichloroacetic acid), OXA (Oxaloacetate), LDH (Lactate dehydrogenase), PDH (Pyruvate dehydrogenase), PDK (Pyruvate dehydrogenase kinase). (**E-F**) HEK293T cells were treated with OXA or DCA at the indicated concentrations for 24 hours. Representative western blots are shown in (E). Quantification of anti-Kla signals relative to lane 1 (untreatment) and normalized to ponceau S staining are shown in (F). Data are represented as mean +/- SEM of three independently performed experiments and each symbol represents an individual experiment. Statistical differences were calculated by 1-way ANOVA followed by Dunnett's correction for multiple comparisons. (**G-I**) Bone-marrow derived macrophages (BMDM) were stimulated with LPS (1 μg/mL) with or without butyrate (2 mM) for 24 hours. Representative western blots are shown in (G). Quantification of anti-Kla and Kac signals relative to lane 1 (untreated, unTx) were shown in

(H) and (I), respectively. Signals were normalized to ponceau S staining. Data are represented as mean +/- SEM of three biological replicates (from three different mice) performed in a single experiment and each symbol represents an individual mouse-derived samples. Statistical differences were calculated by 1-way ANOVA followed by Tukey's correction for multiple comparisons. *$p<0.05$, **$p<0.01$, ***$p<0.001$, ****$p<0.0001$
